## Supplementary figures and images for "Thermodynamic constraints on the assembly and diversity of microbial ecosystems are different near to and far from equilibrium"

### Supplemental Figure 1

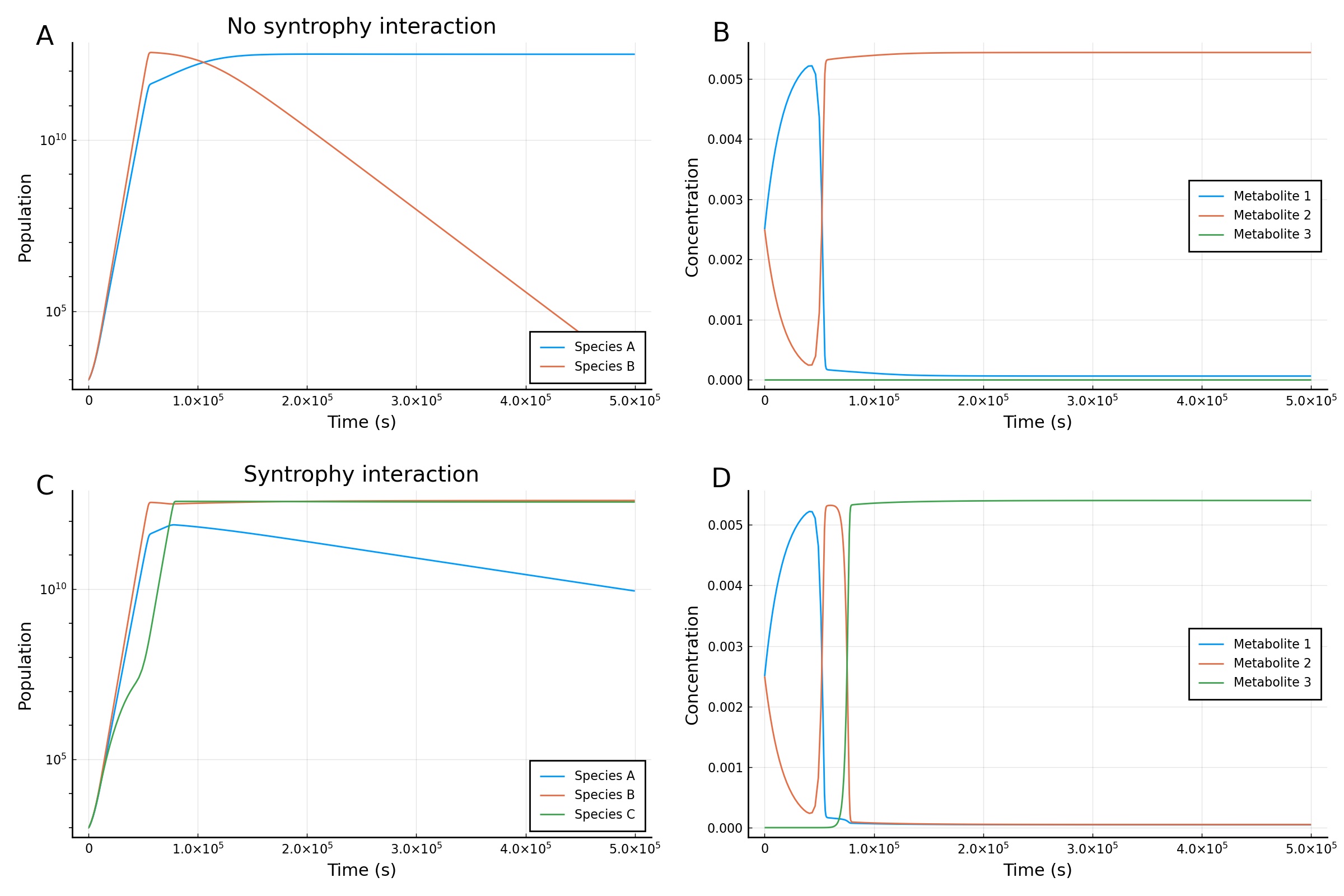

### Supplemental Figure 2

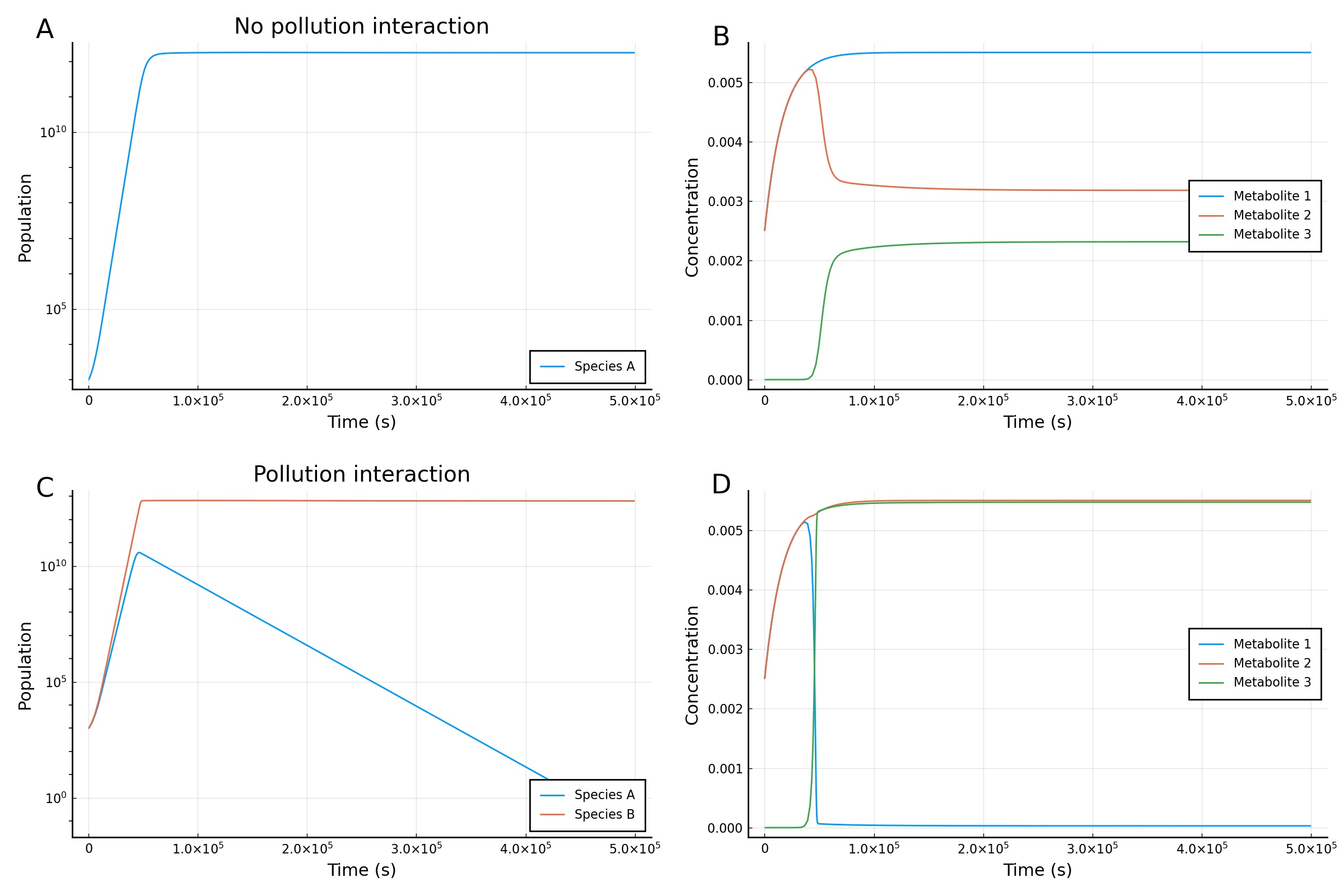

### Supplemental Figure 3

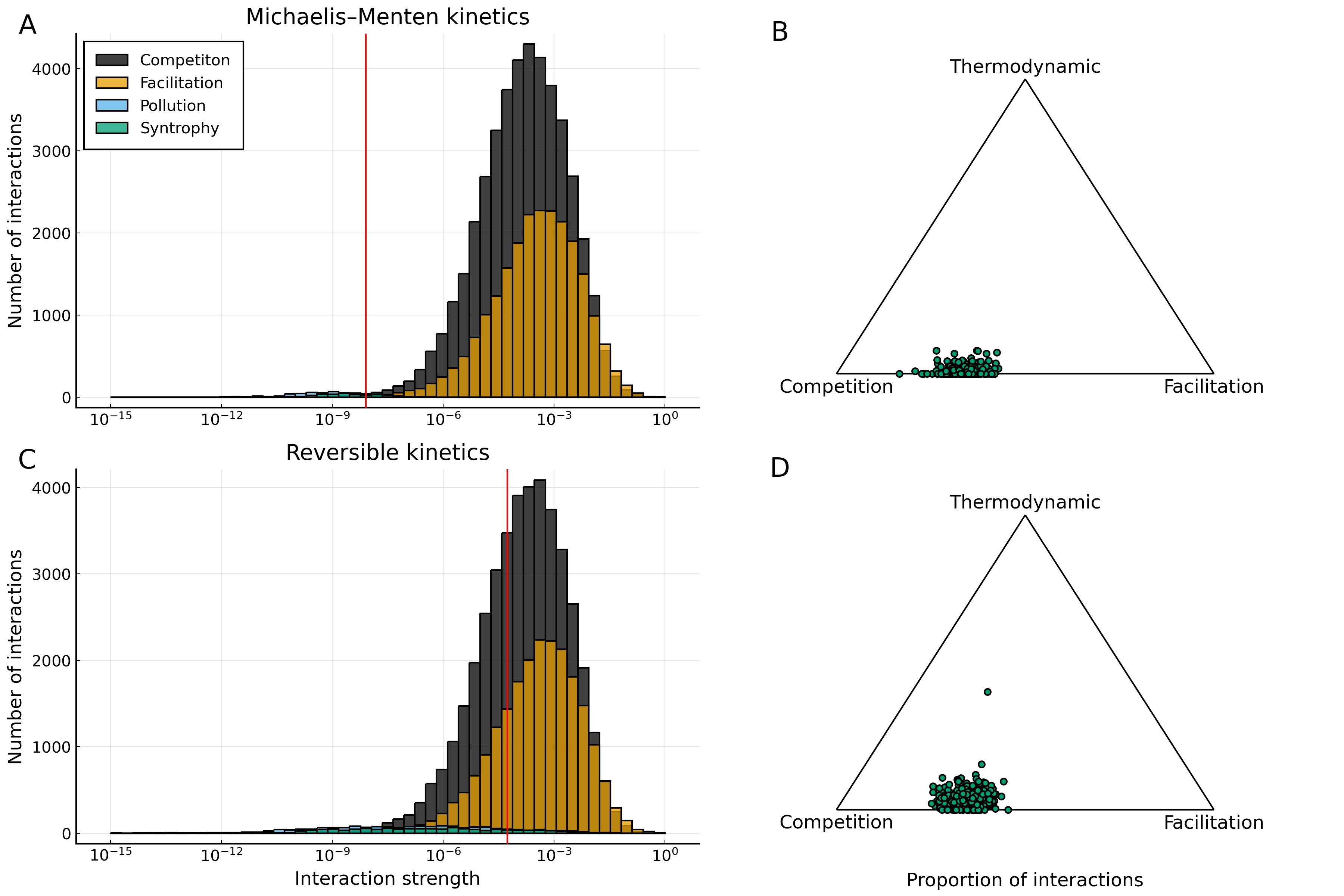
