## Supplemental Text for "Thermodynamic constraints on the assembly and diversity of microbial ecosystems are different near to and far from equilibrium"

**S1 Irreversible Michaelis–Menten kinetics** 2

**S2 Reversible enzyme kinetics** 2

**S3 Proteome partitioning model** 4
S3.1 Derivation of simplified proteome partitioning model . . . . . . . . . . . . . . . . . . 5
S3.2 Validation for single population . . . . . . . . . . . . . . . . . . . . . . . . . . . . . . 7

**S4 Measure of reaction efficiency** 9

**S5 Determining interaction types and strengths** 9

**S6 Comparison with other microbial consumer-resource models** 11

**S7 Interactions between functional groups** 14

**S8 Model parameters** 17

**S9 Analysis of assumptions and parameterisations underlying our model** 17

### S1 Irreversible Michaelis–Menten kinetics

Most models of microbial growth assume irreversible Michaelis–Menten kinetics, which can be represented as

$$E + S \underset{k_{-1}}{\overset{k_{+1}}{\rightleftharpoons}} ES \xrightarrow{k_{\text{cat}}} E + W, \tag{S1}$$

where $k_{+1}$, $k_{-1}$ and $k_{\text{cat}}$ are rate constants, $E$ is an enzyme, $ES$ is an enzyme substrate complex, $S$ is a substrate and $W$ is a waste product. This scheme leads to an expression for the rate of substrate consumption (at steady state) as

$$\frac{d[S]}{dt} = -\frac{d[W]}{dt} = -V_{\text{max}} \frac{[S]}{K_M + [S]},$$

where $V_{\text{max}}$ is the maximal rate of substrate consumption (or waste product production), and $K_M$ is the Michaelis (saturation) constant. These constants are defined in terms of rate constants by

$$V_{\text{max}} = k_{\text{cat}}[E]_0$$

and

$$K_M = \frac{k_{-1} + k_{\text{cat}}}{k_{+1}},$$

where $[E]_0$ is the initial enzyme concentration. This provides a mechanistic justification for the phenomenological Monod microbial growth equation, which is frequently used in ecological modelling [1]. However, the enzyme scheme in Eq. S1 assumes that the second reaction is irreversible which is only possible (as an approximation) if this reaction is dissipating a very large amount of free energy. Though this is a reasonable assumption for cases of growth on high free energy substrate, it is not when considering the low free-energy change reactions commonly observed in microbial ecosystems. We thus wish to consider a fully reversible scheme of enzyme kinetics.

### S2 Reversible enzyme kinetics

To model non-equilibrium thermodynamics, we can instead consider the scheme

$$E + S \underset{k_{-1}}{\overset{k_{+1}}{\rightleftharpoons}} EX \underset{k_{-2}}{\overset{k_{+2}}{\rightleftharpoons}} E + W, \tag{S2}$$

where $k_{\pm 1,2}$ are rate constants [2]. This scheme is fully reversible and is thus suitable to describe low free-energy change reactions. In this case the rate of change of substrate/waste-product concentration (at steady state) can be found to be

$$\frac{d[W]}{dt} = -\frac{d[S]}{dt} = \frac{\frac{V_S}{K_S}[S] - \frac{V_W}{K_W}[W]}{1 + \frac{[S]}{K_S} + \frac{[W]}{K_W}}, \tag{S3}$$

where $V_W$ and $V_S$ are maximal rates for waste product and substrate production, respectively. These rates are defined for the case where all enzymes are bound as complexes ($[EX] = [E]_0$) and can be written as

$$V_S = k_{+2}[E]_0, \qquad V_W = k_{-1}[E]_0.$$

Additionally, $K_S$ and $K_W$ are the dissociation constants for substrate and waste product, respectively. These constants can be defined in terms of rate constants as

$$K_S = \frac{k_{-1} + k_{+2}}{k_{+1}}, \qquad K_W = \frac{k_{-1} + k_{+2}}{k_{-2}}.$$

In order to further simplify Eq. S3 we now make use of a quantity termed the equilibrium constant. This is a thermodynamic constant specific to physical conditions (e.g. temperature, particular reaction type), and constrains the relative values of the rate constants (at a given temperature) by [3]

$$\mathcal{K} = \frac{k_{+1}k_{+2}}{k_{-1}k_{-2}}. \tag{S4}$$

Eq. S3 can then be rearranged to

$$\frac{d[W]}{dt} = \frac{V_S\left([S] - \frac{[W]}{\mathcal{K}}\right)}{K_S + [S] + \frac{K_S}{K_W}[W]}. \tag{S5}$$

The value of the equilibrium constant $\mathcal{K}$ will be affected by the total Gibbs free energy ($\Delta G_T$) of the reaction and the endergonic reactions it is coupled to. Coupling one or more endergonic reactions ($\Delta_r G > 0$) to our original exergonic reaction ($\Delta_r G < 0$) allows the acquisition of free energy for later use in other processes. Cells commonly couple metabolic reactions to the production of ATP which is strongly endergonic in order to transduce free energy [4]. The coupling of our generic reaction to the synthesis of ATP can be expressed as

$$E + S + \eta\text{ADP} + \eta P_i \underset{k_{-1}}{\overset{k_{+1}}{\rightleftharpoons}} EX \underset{k_{-2}}{\overset{k_{+2}}{\rightleftharpoons}} E + W + \eta\text{ATP},$$

where $P_i$ is organic phosphate, $\eta$ is the number ATP produced per reaction event. In order to account for alternative free energy transducing processes such as the pumping of ions across membranes, $\eta$ can to take non-integer values in our full model [5]. The total Gibbs free energy of a reaction coupled to ATP production in this manner is

$$\Delta G_T = \Delta_r G^0 + RT \ln Q + \eta \Delta G_{\text{ATP}}, \tag{S6}$$

where $R$ is the gas constant, $T$ is the temperature, $\Delta_r G^0$ is the Gibbs free energy of one mole of reaction at standard conditions, $Q$ is the reaction quotient and $\Delta G_{\text{ATP}}$ is the Gibbs free energy per mole of ATP. The value of $\Delta G_{\text{ATP}}$ is dependant on the cellular concentrations of ATP, ADP and organic phosphate. In this work we will assume that these are constant and will use the experimentally-derived estimate for $\Delta G_{\text{ATP}}$ of 75 kJ mol$^{-1}$ throughout [6]. For the equilibrium case $\Delta G_T = 0$ the reaction quotient $Q$ takes the value

$$Q_0 = \mathcal{K} = \exp\left(\frac{-\Delta_r G^0 - \eta \Delta G_{\text{ATP}}}{RT}\right),$$

equal to the previously-discussed equilibrium constant. This means that the constraint on the relative reaction rates (shown in Eq. S4) is determined by the reaction quotient of the combined reaction at equilibrium. The first term in the numerator is the Gibbs free energy change at standard conditions (i.e. the substrate and waste product have equal concentration). The equilibrium constant thus tells in which direction we would expect the reaction to proceed when waste product and substrate have equal concentrations (i.e. towards increasing waste product if $\mathcal{K} > 1$ and towards increasing substrate if $\mathcal{K} < 1$). The second term of the numerator shifts this equilibrium so as the amount of free energy dissipated decreases the amount of waste product that needs to build up for equilibrium to be reached also decreases. Increasing temperature $T$ decreases the magnitude of this exponent and so the effect varies dependant on the sign of the numerator. In our model we will neglect the effect of temperature on the dynamics.

Following [3], we can make use of the fact that $V_S = k_{+2}[E]_0$ (which we shall simplify to $k[E]_0$) to further rearrange Eq. S5 to

$$\frac{d[W]}{dt} = \frac{k[E]_0\left([S] - \frac{[W]}{\mathcal{K}}\right)}{K_S + [S] + \frac{K_S}{K_W}[W]} = \frac{k[E]_0[S]\left(1 - \frac{[W]}{[S]\mathcal{K}}\right)}{K_S + [S]\left(1 + \frac{K_S[W]}{K_W[S]}\right)}.$$

Into to the above expression we can substitute $\frac{K_S}{K_W} = \frac{r}{\mathcal{K}}$ and $\theta = \frac{Q}{\mathcal{K}}$, and then obtain the consumption rate $q$ as

$$q([S],[W]) = \frac{k[E]_0[S](1-\theta)}{K_S + [S](1+r\theta)}, \tag{S7}$$

where $k$ is the maximal reaction rate, $K_S$ is the enzyme (substrate) saturation constant, and $r$ is a reversibility factor [3]. Further, $\theta$ is a thermodynamic factor that tends towards one when reactions are close to equilibrium, which can be expressed as

$$\theta = \theta\left([S],[W]\right) = \frac{Q([S],[W])}{\mathcal{K}(S,W)}$$

where $\mathcal{K}(S,W)$ is the equilibrium constant for the reaction from $S$ to $W$ and $Q$ is the reaction quotient given by

$$Q(A,B) = \frac{B}{A}.$$

It is useful to note that for reactions dissipating large amounts of free energy $\mathcal{K} \gg Q$, and thus $\theta \approx 0$, meaning that Michaelis–Menten dynamics are recovered from Eq. S7. This explains why the Michaelis–Menten scheme is appropriate to model many (but not all) ecosystems, particularly lab cultures where free energy supply is generally high.

#### S3 Proteome partitioning model

The basic assumption of our proteome partitioning model is that consistent with empirical observations, cellular proteins can be divided into three broad classes: 1) ribosomes (denoted $R$) that allow further protein synthesis, 2) metabolic enzymes (denoted $P$) that obtain energy from the external metabolites, and 3) a fixed proportion of housekeeping proteins (denoted $Q$) necessary for other vital cellular processes [1]. The rate of change of these three classes of proteins can be modelled with the following equations:

$$\begin{aligned}
\frac{dR}{dt} &= \frac{\gamma(a)}{n_R} f_b^R R - \lambda R \\
\frac{dP}{dt} &= \frac{\gamma(a)}{n_P} f_b^P R - \lambda P \\
\frac{dQ}{dt} &= \frac{\gamma(a)}{n_Q} f_b^Q R - \lambda Q.
\end{aligned}$$

Here, $\lambda$ is the cellular growth rate, $a$ is the cellular energy concentration, $\gamma(a)$ is the effective translation elongation rate, $n_x$ is the typical number of translation steps needed to produce a protein of type $x$, and $f_b^x$ is the fraction of ribosomes bound and producing protein of type $x$. Further, the number of translations steps needed for each protein step can be approximated from the literature as 7459 (amino acids per protein) [7], 300 and 300 [8, 9] for $n_R$, $n_P$ and $n_Q$, respectively. As protein synthesis operates far from thermodynamic equilibrium in order to minimise translation errors, we assume it to be effectively irreversible. The effective translation elongation rate thus has a saturating dependence on energy availability as

$$\gamma(a) = \frac{\gamma_m a}{\gamma_{\frac{1}{2}} + a},$$

where $\gamma_m$ is the maximum elongation rate and $\gamma_{\frac{1}{2}}$ is the free energy concentration where the rate is half of the maximum. To the best of our knowledge, this half-maximum constant has not been empirically measured, so is a free parameter in our model. However, the maximum elongation rate ($\gamma_m$) has been measured to be 21.0 amino acids per second [10].

#### S3.1 Derivation of simplified proteome partitioning model

The dynamics for the total mass of the cell (in amino acids) are given by

$$\frac{dm}{dt} = \gamma(a) f_b R - \lambda m \tag{S8}$$

where $f_b$ is the total fraction of ribosomes that are bound. We assume that the cells retain a constant average protein mass, which an order of magnitude estimate of is taken from the literature as $m = 10^8$ [10]. As $m$ is taken to be constant at steady state, Eq. S8 must equal zero and so can be solved to give an expression for growth rate as

$$\lambda = \frac{\gamma(a) f_b R}{m}. \tag{S9}$$

Although the fraction of ribosomes bound ($f_b$) would be expected to vary with the cells internal energy concentration, as a simplifying assumption we take it to be fixed at a value of 0.7 [11]. However, the fractions of ribosomes bound making specific protein classes ($f_b^R$ etc) can still vary. Previous mechanistic modelling in the literature suggests that the transcription rate of ribosomal mRNAs shows a stronger response to increasing energy concentration than the transcription rate of non-ribosomal mRNAs [9]. This means that as the cell's energy concentration increases the relative proportion of ribosomes bound to ribosomal mRNAs, i.e. $f_b^R$ increases at the expense of $f_b^P$ as $f_b^Q$ is held constant by steep auto-inhibition. Thus, the fraction of the proteome dedicated to each protein type should be expected to be approximately a direct function of the energy concentration. Each protein class can be expressed as a fraction of the total proteome satisfying the following constraint

$$\phi_R + \phi_P + \phi_Q = 1,$$

where $\phi_R = f_b^R/f_b$ is the ribosome fraction, $\phi_P = f_b^P/f_b$ is the metabolic enzyme fraction, and $\phi_Q = f_b^Q/f_b$ is the housekeeping fraction. The maximum value that the ribosome fraction can reach is $\phi_R^{\max} = 1 - \phi_Q$, the cells response to increasing energy concentration must therefore be saturating. As we have no specific mechanism to link this response function to, we assume that it is of Michaelis–Menten form. We can express the ideal ribosome fraction $\phi_R^*$ (main text Eq. 7) as a function of energy concentration $a$ by

$$\phi_R^*(a) = \frac{a}{a + \Omega_{\frac{1}{2}}} (1 - \phi_Q) \tag{S10}$$

where $\Omega_{\frac{1}{2}}$ is the half-maximum constant with respect to energy concentration and $\phi_Q = 0.45$ [12]. As the response function does not correspond to a specific mechanism, $\Omega_{\frac{1}{2}}$ is a treated free parameter of the model. The form of Eq. S10 ensures that as the energy concentration approaches infinity the ideal ribosome fraction approaches its maximum possible value of $1 - \phi_Q$ and as the energy concentration approaches zero the ideal metabolic fraction $\phi_P^*$ approaches this same maximum.

This ideal proteome partition cannot be immediately realised by the cell, as the synthesis of large quantities of protein is required in order to shift the relative size of the fractions. The actual ribosome fraction $\phi_R$ dynamics (Eq. 3 main text) are thus described by

$$\frac{d\phi_R}{dt} = \frac{1}{\tau_g}\left(\phi_R^*(a) - \phi_R\right) \tag{S11}$$

where $\tau_g$ is the characteristic time scale of proteome shifting. In order to define the characteristic time scale we consider the extreme (and unrealisable) case where the ideal ribosome fraction is equal to the maximum $(1 - \phi_Q)$ but the actual fraction is zero. We assume that in this extreme case only metabolic proteins would be synthesised, so that each biomass doubling would halve metabolic protein fraction. The ideal fraction will only ever be asymptotically approached through this process of halving, so we consider the process to be complete when the metabolic fraction has dropped to 1% of the maximum. The characteristic time scale can therefore be expressed as

$$\tau_g = \frac{1}{\lambda}\frac{\log 100}{\log 2}, \tag{S12}$$

where $\lambda^{-1}$ is the time required for biomass doubling and $\frac{\log 100}{\log 2}$ is the number of halvings required to go from 100% to 1%. In cases where the ideal ribosome fraction is less extreme we would expect the protein production to be balanced between both types. This means that the reduction in time due to needing to synthesise less protein to reach the ideal fraction would be at least partially counteracted by a lower rate of synthesis of the deficient protein type. Thus, we expect the characteristic timescale ($\tau_g$) to be relevant in all cases.

Growth rate (Eq. S9) can now be re-expressed in terms of the protein fraction as

$$\lambda = \frac{\gamma(a) f_b}{m}\frac{m}{n_R}\phi_R = \frac{\gamma(a) f_b \phi_R}{n_R}, \tag{S13}$$

which is Eq. 5 in the main text. The growth rate is thus heavily determined by the internal energy concentration. The full dynamics of the cells energy concentration is given by

$$\frac{da}{dt} = J - \mathcal{T} - \lambda a, \tag{S14}$$

where $J$ is the rate at which the cell acquires free energy (as ATP) and $\mathcal{T}$ is the rate of free energy consumption due to protein translation. Following [9], we here make the assumption that protein translation is the only significantly free energy consuming process in the cell. The rate of ATP consumption by protein translation can be expressed as

$$\mathcal{T} = \chi m \lambda,$$

where $\chi$ is the ATP consumption per translation step, and $m\lambda$ gives the number of translation steps per unit time. The value of $\chi$, including the cost of amino acid synthesis, is taken to be 29.0 [13]. In order to find an expression for the energy acquisition rate $J$ we must further divide the proteome. The metabolic protein fraction is in turn divided between a multiple different free energy yielding reactions subject to the constraints

$$\sum_{\alpha \in O} \nu_\alpha = 1 \tag{S15}$$

and

$$\nu_\alpha \geq 0 \quad \forall \alpha, \tag{S16}$$

where $\nu_\alpha$ is the relative expression of reaction $\alpha$ and $O$ is the set of reactions possessed by the cell. The amount of a enzyme available for reaction $\alpha$ can then be expressed as

$$E_\alpha = \frac{m\nu_\alpha \phi_P}{n_P},$$

where $n_P$ is the mass (i.e. number of translation steps) of a metabolic protein. The rate of free energy acquisition can now be expressed as

$$J = \sum_{\alpha \in O} \eta_\alpha q_\alpha(E_\alpha, S_\alpha, W_\alpha), \tag{S17}$$

where $\eta_\alpha$ is the amount of ATP the cell synthesises from reaction $\alpha$. The reaction rate for reaction $\alpha$ is a slightly altered form of Eq. S7

$$q_\alpha(E_\alpha, S_\alpha, W_\alpha) = \frac{k_\alpha E_\alpha S_\alpha (1-\theta)}{K_{S_\alpha} + S_\alpha(1 + r_\alpha \theta)},$$

where $k_\alpha$ is the rate limiting reaction rate ($k_{+2}$ in Eq. S2) for reaction $\alpha$, $K_{S_\alpha}$ the saturation constant, and $r_\alpha$ is the reversibility factor.

The cell population is denoted by $N$ and the population dynamics can be described by

$$\frac{dN}{dt} = N\left(\lambda - d\right),$$

where $d$ is the death rate accounting for both explicit cell death and other biomass losses. Our model incorporates a large number of metabolites, which can perform the function of both substrates and products for the various reactions. The concentration of metabolite $\beta$ is given by $C_\beta$, and its dynamics can be described by

$$\frac{dC_\beta}{dt} = \kappa_\beta - \rho_\beta C_\beta + \left(p_\beta(\mathbf{C}) - c_\beta(\mathbf{C})\right) N, \tag{S18}$$

where $\kappa_\beta$ is the supply rate of metabolite $\beta$, $\rho_\beta$ is the dilution rate of metabolite $\beta$, $p_\beta$ is the rate at which the cells produce metabolite $\beta$, and $c_\beta$ is the rate at which the cells consume metabolite $\beta$. These net reaction rates can be expressed as

$$p_\beta(\mathbf{C}) = \sum_{\alpha \in O} \delta_{C_\beta, W_\alpha} q_\alpha(E_\alpha, S_\alpha, W_\alpha),$$

and

$$c_\beta(\mathbf{C}) = \sum_{\alpha \in O} \delta_{C_\beta, S_\alpha} q_\alpha(E_\alpha, S_\alpha, W_\alpha).$$

where $\delta_{C_\beta, W_\alpha}$ and $\delta_{C_\beta, S_\alpha}$ are Kronecker deltas that are zero unless metabolite $\beta$ is the waste product of reaction $\alpha$ or metabolite $\beta$ is the substrate of reaction $\alpha$, respectively.

#### S3.2 Validation for single population

In order to establish the validity of our model we attempted to replicate the linear growth laws found in [1]. To do this we simulated the strains with the substrate and waste product concentrations held fixed. We then plotted the final ribosome fraction $\phi_R$ against the final growth rate $\lambda$ (due to the fixed nutrient environment the strains exponentially grow forever). The first growth law that we consider is found in experiments where drugs decreasing the rate of protein translation are applied to an exponentially growing population. In these experiments as growth rate decreases a linear increase in ribosome fraction is observed. To approximately replicate these experiments in our model we decreased the maximum translation rate per ribosome $\gamma_m$ (Fig. ST1, top). The other growth law that we consider is found in experiments where microbial strains are grown upon media of successively higher nutrient quality. Here, as growth rate increases ribosome fraction linearly increases along with it. To adapt our framework to this experimental setup we used $\Delta_r G^0$ is the standard Gibbs free energy of the reaction as a proxy for nutrient quality. As the magnitude of this Gibbs free energy is increased there is more free energy available per reaction event to be retained as ATP and thus the value of $\eta$ can be increased (Fig. ST1, bottom).

We found that across most growth rates linear growth laws could be recovered (see Fig. ST1). However, for both growth laws at low growth rates deviations were observed. In the case of the nutrient quality law at low growth rates the ribosome fraction is lower than would be expected based on the growth rate. This arises because when strains utilise low free energy substrates significantly less ATP accumulation occurs. This acts to further reduce the growth rate. Hence, as nutrient quality is increased the growth rate initially increases super-linearly. In the case of the translational inhibition law at low growth rates the ribosome fraction is higher than would be expected based on the growth rate. In this case the ability of the strain to make use of energy is significantly hampered by the decreased maximum translation rate $\gamma_m$, causing accumulation of energy by the cell. This energy accumulation partially compensates for the decreased maximum translation rate, so growth rate declines more slowly than would be expected.

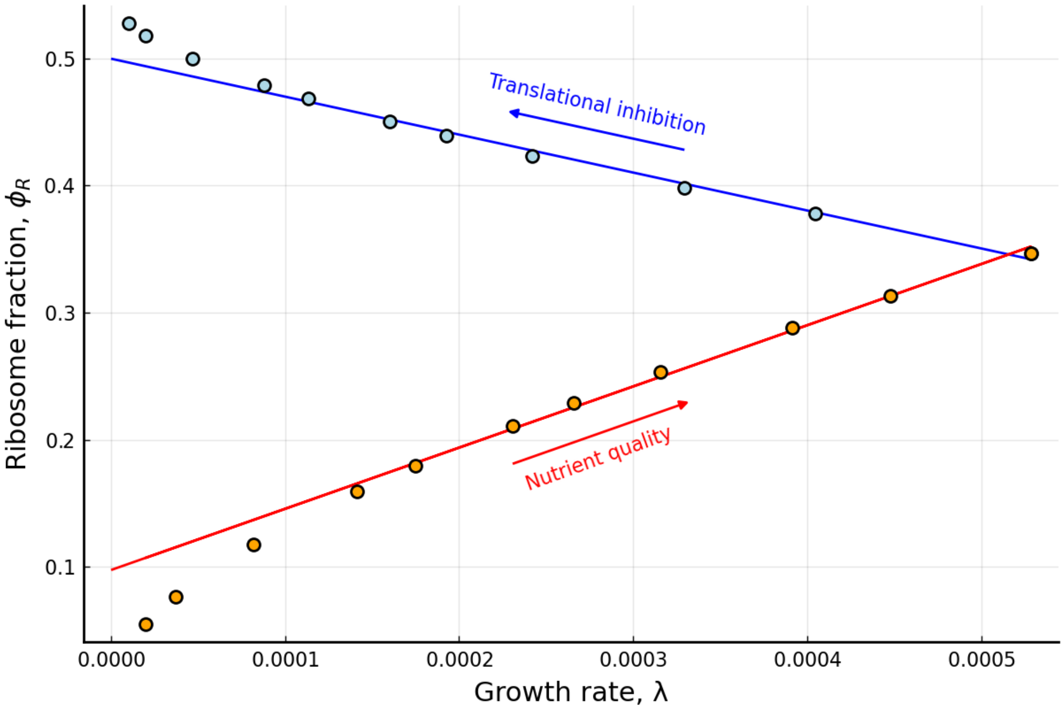

Figure ST1: **Validation based on growth laws.** Final ribosome fraction is plotted against final growth rate for our model simulated using various conditions. The upper (blue) data are simulated for decreasing maximum translation rate $\gamma_m$. The lower (orange) data are simulated for increasing standard reaction Gibbs free energy (a proxy for nutrient quality). For both sets of data the linear fit is found excluding the three points with the lowest growth rate.

#### S4 Measure of reaction efficiency

In order to assess how the overall (thermodynamic) efficiency of the community changes through the assembly process, we need to calculate the efficiency of each reaction for each species. The free energy change of a reaction is not fixed as it depends on both the substrate and waste-product concentrations (see Eq S6). We measure the efficiency against the maximum possible Gibbs free energy transduction, which is determined using the minimum equilibrium product to substrate ratio allowed when choosing $\eta$ values ($s_r = 10^{-2}$, detailed in the Simulations section of the main text). Species $i$'s efficiency for reaction $\alpha$ is thus given by

$$\zeta_{\alpha,i} = \frac{-\eta_{\alpha,i}\Delta G_{\mathrm{ATP}}}{\Delta_\alpha G^0 + RT\ln(s_r)},$$

where $\eta_{\alpha,i}$ is the number of ATP generated per reaction event by species $i$ for reaction $\alpha$, $\Delta G_{\mathrm{ATP}}$ is the free energy per mole of ATP, $\Delta_\alpha G^0$ is reaction $\alpha$'s free energy change (per mole), $R$ is the gas constant, and $T$ is the temperature. It is worth noting that the free-energy per mole of ATP varies substantially with temperature. However, throughout this work we consider a fixed temperature and so assuming the free-energy of ATP hydrolysis to be constant is a reasonable assumption. For each simulation the average value of reaction efficiency across all reactions for all species can then be calculated by

$$\bar{\zeta} = \sum_{i=1}^{B} N_i \left( \sum_{\alpha \in O_i} \nu_{\alpha,i}\zeta_{\alpha,i} \right), \tag{S19}$$

where $B$ is the number of species surviving in the simulation, $N_i$ is the $i^{\mathrm{th}}$ species' population abundance, $O_i$ is the set of reactions possessed by species $i$, and $\nu_{\alpha,i}$ is the relative expression of reaction $\alpha$ by species $i$. The average reaction efficiency shown in Fig 4D (main text) is calculated using Eq S19.

#### S5 Determining interaction types and strengths

All interactions in our model occur via microbes consuming or producing metabolites. Thus, inferring interactions between strains involves calculating both the responses of individual species' populations to changes in metabolite concentrations, and the reciprocal impact on the metabolite concentrations. Calculating these 'susceptibilities' analytically can allow solutions that link interactions to emergent behaviours of such systems. An example is the recent work applying the cavity calculation of statistical physics to model ecosystems [14, 15], which allowed susceptibilities to be linked to the ability of new species to invade. However, analytical solutions for the effective resource capacity and the resource susceptibility could not be found in our model, as reaction rates depend not just on substrate concentrations but also waste-product concentrations. This generates a coupling between resource dynamics (i.e. where the concentration of one metabolite can effect the rate a different metabolite is consumed at), which makes the system far harder to solve analytically. In the absence of the previously mentioned analytic results performing a calculation reminiscent of the cavity calculation does not seem feasible. So rather than making use of this elegant method of determining whether newly introduced species can persist we will make use of our susceptibilities in a relatively crude manner to calculate the instantaneous impact of each species on every other species at steady state, as a measure of interactions between species. These interactions can only be accurately calculated when the ecosystem has reached steady state.

So, in order to calculate species' responses we consider a perturbation of a single metabolite ($\beta$) that takes the form

$$\mathbf{C'}_\beta = [C_1, \ldots, (1+\xi) \times C_\beta, \ldots, C_M],$$

where $C_\beta$ is the steady state value of metabolite $\beta$, and $\xi$ is the (fractional) perturbation size. We then determine the effect this perturbation has on the rate of change of ATP (Eq. S14) for strain $i$ which is described by

$$\frac{da_i}{dt}\left(\mathbf{C'}_\beta\right) = J_i\left(\mathbf{C'}_\beta\right) - \chi m \lambda_i - a_i \lambda_i, \tag{S20}$$

where $J_i$ is the rate of ATP generation (Eq. S17), $\chi$ is the ATP consumption per translation step, and $m$ is the total mass of the cell (in amino acids). The second term of Eq. S20 gives the rate at which the strain uses ATP to synthesise protein, and the third term gives the rate at which ATP is diluted by volume expansion. The metabolite perturbation only initially effects the ATP generation rate $J_i$, as the other two terms have no direct dependence on metabolite concentrations. We cannot find the effect that the perturbation has on the growth rate (Eq. S13) directly without finding the new steady state value of $a_i$, which can only be found numerically. Therefore, we instead assume that the change in the (per cell) rate of ATP generation ($J_i$) directly impacts the rate of change of (population) biomass abundance as

$$\frac{dN_i}{dt}\left(\mathbf{C'}_\beta\right) = \frac{N_i}{\epsilon}\frac{dJ_i}{dt}\left(\mathbf{C'}_\beta\right) = \frac{N_i}{\epsilon}\frac{da_i}{dt}\left(\mathbf{C'}_\beta\right),$$

where $\epsilon$ is the cost (in ATP units) to synthesise one cell. By assuming that the dilution ($3^{\text{rd}}$) term in Eq. S20 is sufficiently small as to be neglected this cost simply becomes $\epsilon = \chi m$, and the rate of change of biomass in response to the metabolite perturbation can then be found as

$$\frac{dN_i}{dt}\left(\mathbf{C'}_\beta\right) = \frac{N_i}{\chi m}\frac{da_i}{dt}\left(\mathbf{C'}_\beta\right). \tag{S21}$$

This allows us to quantify the impact of the perturbation of a specific metabolite $\beta$ on each strain's population abundance.

We also need to find the impact the specific strains have on the metabolite concentrations. We do this by finding the steady state nutrient concentrations, which for multiple strains are described by an extension of Eq. S18 as

$$\frac{dC_\beta}{dt} = \kappa_\beta - \rho_\beta C_\beta + \sum_{i=1}^{B}\left(p_{i,\beta}(\mathbf{C}) - c_{i,\beta}(\mathbf{C})\right)N_i = 0,$$

where $B$ is the total number of strains. If consumption ($c$) and production ($p$) rates are known the steady state value of metabolite $\beta$ can be found as

$$C^*_\beta = \frac{\kappa_\beta + \sum_{i=1}^{B}\left(p_{i,\beta}(\mathbf{C}) - c_{i,\beta}(\mathbf{C})\right)N_i}{\rho_\beta}. \tag{S22}$$

The instantaneous effect of a population perturbation on the concentration will occur due to the changed strain abundance, and the production and consumption rates will only change once the metabolite concentrations have begun to change in response to this. We therefore make the simplifying assumption that the production and consumption rates remain constant. We now calculate the steady-state metabolite concentration (under this assumption) as a proxy for the strength of the instantaneous response as

$$C^*_\beta + \Delta C_{j,\beta} = \frac{\kappa_\beta + \sum_{i\neq j}\left(p_{i,\beta}(\mathbf{C}) - c_{i,\beta}(\mathbf{C})\right)N_i + \left(p_{j,\beta}(\mathbf{C}) - c_{j,\beta}(\mathbf{C})\right)\left(N_j + \Delta N_j\right)}{\rho_\beta}, \tag{S23}$$

where $\Delta N_j$ is the population perturbation of strain $j$ and $\Delta C_{j,\beta}$ is the resulting change in the steady-state value of metabolite $\beta$. By factoring out Eq. S22 from Eq. S23 the size of this change is found to be

$$\Delta C_{j,\beta} = \frac{\left(p_{j,\beta}(\mathbf{C}) - c_{j,\beta}(\mathbf{C})\right)\Delta N_j}{\rho_\beta}. \tag{S24}$$

This concentration change will not be realised (as production and consumption rates adapt), but instead stands as a proxy for the strength of the metabolite's instantaneous response to the population perturbation.

The impact of one strain ($j$) upon another strain ($i$) can now be determined using the two metrics derived in Eqs. S21 & S24. If the population response of strain $i$ (Eq. S21) is positive for a positive perturbation ($\xi > 0$) the strain either solely or predominately uses the metabolite as a substrate to grow. Whereas if the response is negative, the strain is predominately inhibited by the metabolite as a waste product. When a positive perturbation of strain $j$'s population ($\Delta N_j > 0$) results in a positive concentration change for metabolite $\beta$ (Eq. S24) the strain is a net producer, and if it's negative the strain is a net consumer. This behaviour can be used to classify four interaction types. They are 'competition' ($\Delta C_{j,\beta} < 0$ and $\frac{dN_i}{dt}(\mathbf{C'}_\beta) > 0$), 'facilitation' ($\Delta C_{j,\beta} > 0$ and $\frac{dN_i}{dt}(\mathbf{C'}_\beta) > 0$), 'syntrophy' ($\Delta C_{j,\beta} < 0$ and $\frac{dN_i}{dt}(\mathbf{C'}_\beta) < 0$), and 'pollution' ($\Delta C_{j,\beta} > 0$ and $\frac{dN_i}{dt}(\mathbf{C'}_\beta) < 0$). The strength of the impact of strain $j$ upon strain $i$ via metabolite $\beta$ is given by

$$\left(\frac{dN_i}{dt}\right)_{j,\beta} = \frac{dN_i}{dt}(\mathbf{C'}_\beta)\frac{1}{\xi C_\beta}\Delta C_{j,\beta}$$

where the $\xi C_\beta$ term converts the concentration change ($\Delta C_{j,\beta}$) into units of the metabolite perturbation size ($|\mathbf{C'}_\beta|$). A positive interaction type implies a facilitation or syntrophy interaction, whereas a negative interaction strengths implies a competition or pollution interaction. The interaction strengths obtained from this expression will not be symmetric, i.e. $|(\frac{dN_i}{dt})_{j,\beta}| \neq |(\frac{dN_j}{dt})_{i,\beta}|$. Additionally, having an interaction type between strain $i$ and strain $j$ does not necessarily determine the type of the converse interaction. Competition interactions are always paired, but pollution interactions only pair if both strains are sufficiently close to thermodynamic equilibrium, and facilitation only pairs with syntrophy when the facilitating strain is close to equilibrium. The total strength of interaction from strain $j$ upon strain $i$ can be calculated by summing over the metabolites as

$$\left(\frac{dN_i}{dt}\right)_j = \sum_{\beta=1}^{M}\left(\frac{dN_i}{dt}\right)_{j,\beta},$$

where $M$ is the total number of metabolites. This total interaction strength is not used in the main text as it doesn't allow for a full classification of interaction types.

### S6 Comparison with other microbial consumer-resource models

In this section, we compare our model with previous theoretical work in the area of microbial consumer-resource models.

#### S6.1 Previous work addressing violations of competitive exclusion theory

In nature, a huge diversity of microbes is observed, despite the expectation from competitive exclusion theory that only one consumer species can survive per limiting resource [16]. This apparent contradiction has been the subject of much historic discussion in the literature (exemplified by Hutchinson's "paradox of the plankton"; [17]). Many potential resolutions to this contradiction have been proposed, such as chaotic dynamics [18], temporal heterogeneity [19], spatial heterogeneity [20], limitation by factors other than resource availability (e.g. predation) [21], or rapid evolution [22]. There are also some approaches that bear strong similarity to ours. The first of these develops a thermodynamic consumer-resource model based on irreversible Michaelis–Menten kinetics [23]. Using this model Großkopf et al. observe that two species can coexist on the same substrate, provided that they use different reactions (with different products) to do so. This represents a solution to the coexistence constraint predicted by the classical competitive exclusion principle, and suggests thermodynamic limitations can help explain the paradox, more broadly. In another approach that bears similarity to ours, a consumer-resource model was developed with a fixed enzyme budget that can be divided between metabolites, all of which can support cell growth and reproduction [24]. Provided that all species have enzyme budgets of the same size and the resource supply rate are contained within the convex hull of metabolic strategies, an indefinite number of species can survive on just three supplied metabolites. It should however be noted (see [25]), that this result only holds if both total enzyme budget and biomass loss rates are fixed between species. Additionally, this model only considers metabolite consumption and does not include the commonly observed production of metabolites by microbes.

The proteomic trade-off that our model includes represents a more detailed version of the previously mentioned metabolic trade-off modeled in previous studies [24, 25]. Our more realistic "enzyme budget" is inversely proportional to the size of the ribosome fraction rather than being fixed. In addition to this, our model includes thermodynamic inhibition (similarly to [23]). It is, therefore, natural to ask whether the results found in the previously discussed approaches are obtained with our model. Specifically, whether the increased diversity is retained over the longer run in larger and more realistic stochastically-generated communities. We observe that the average number of survivors is lower than the average number of available substrates (see Fig 3B main text), and that the number of survivors never exceeds the number of diversified substrates (see Fig ST2). Our results suggest that although metabolic or thermodynamic trade-offs can offer a solution to the mismatch between competitive exclusion theory and ecological reality in special cases, this is not a general effect.

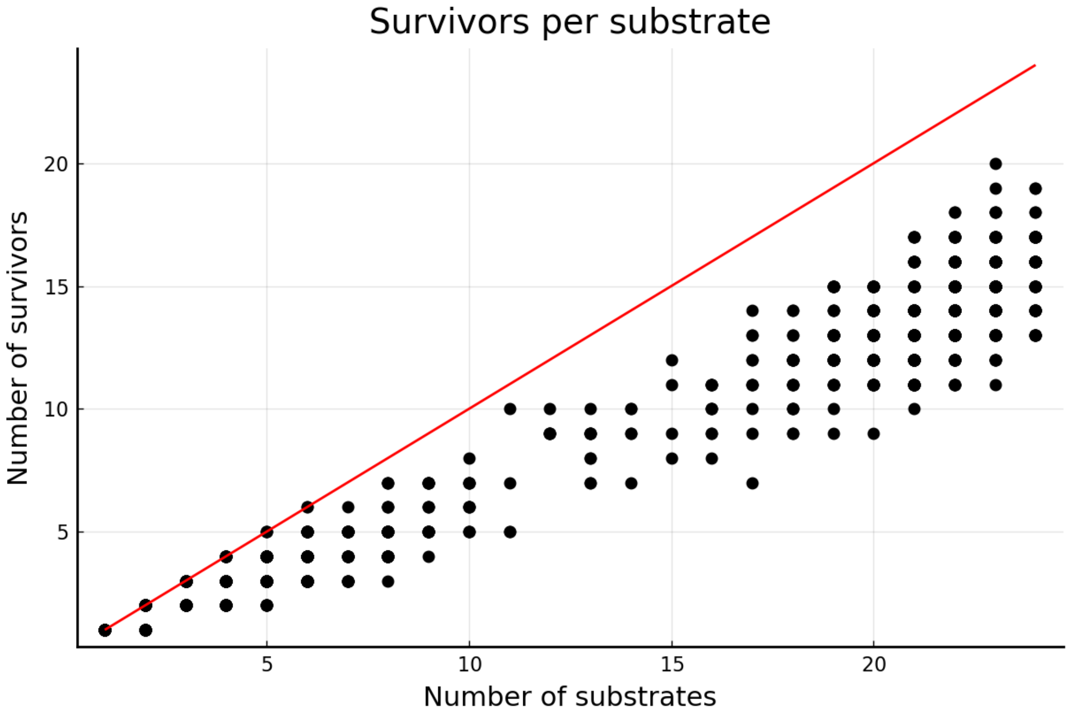

Figure ST2: **Competitive exclusion in our model.** Plot of the number of surviving species against the total number of substrates diversified by the time steady state is reached. This shows the full data from across all four regimes considered in the main text, e.g. 1000 simulation runs in total. In no case does the number of survivors exceed the number of substrates (equality indicated by the red line).

#### S6.2 Previous work modelling adaptive proteome allocation between reactions

In another recent approach, a proteome trade-off was incorporated into a consumer-resource model [26]. In this model, metabolites are consumed but not produced by species, and every species splits its enzyme investment between the different metabolites. This model fundamentally differs from the one presented in Posfai et al. [24], in that the size of the enzyme budget changes with the size of the ribosome fraction, and the fraction of a species' enzyme budget dedicated to each metabolite is updated using an optimisation procedure. As previously mentioned, our model captures the first of these effects, but in contrast, our model treats the proportional reaction expressions as fixed (see Eq 9 main text), meaning that overall reaction expression of the community can change only through species sorting. The inclusion of metabolite production would lead to a significantly more complex and computationally intensive optimisation procedure. We therefore chose to use fixed reaction expressions, in order to keep our model tractable for the study of the assembly of complex microbial communities.

Using their model, Pacciani-Mori et al. [26] find that the number or surviving consumer species can exceed the number of substrates, provided that the biomass loss rate for each species is proportional to the proteome fraction allocated to both metabolic protein and ribosomes (i.e. the non-'housekeeping' fractions). This assumption does not require that different species must have the same biomass loss rate when they possess the same 'housekeeping' fraction, and so is a far less restrictive assumption than the one made by Posfai et al. In our model, all species have the same biomass loss rates and 'housekeeping' fractions so this condition is met. However, in contrast to our model, it is implicitly assumed that all species are equally good competitors for the various substrates (i.e. equal uptake rates per unit protein investment in a particular substrate). Species are therefore selected for their overall metabolic strategies, rather than being intrinsically better competitors for particular substrates. Our model breaks this equality using stochastic fluctuations about a mean value, which could explain why we never observe more surviving species than substrates at steady-state (see Fig ST2). However, this could alternatively be due to the lack of adaptive proteome allocation between different reactions in our model. Determining which of these two is the case would require the development of a model that combines both adaptive reaction expressions and realistic (non-equilibrium) thermodynamic constraints.

#### S6.3 Previous work addressing effect of energy supply on diversity

Recent theoretical work has addressed the link between energy supply and steady state microbial community diversity [27]. Marsland et al.'s model incorporates microbial metabolite production in addition to consumption, with the level of metabolite exchange between species being governed by the leakage parameter. In this model, maximum diversity is limited directly by the total amount of energy leaked by species (as metabolites), which determines effectively how many substrates (metabolites) are available (in sufficient quantities) to sustain a diverse set of species. Therefore, diversity is higher when energy supply is large with a significant proportion of it leaked. In our model, the rate of metabolite leakage scales directly with the reaction rate, therefore their leakage parameter has no obvious equivalent in our framework. Despite this, their result is broadly consistent with ours (Fig 3 main text).

A key difference between our model and theirs, is that they do not introduce a metabolite (free-energy) hierarchy. This means that the consumer-resource dynamics in their model involve relatively short transients because metabolite uptake, allocation to growth and leakage is instantaneous, and any metabolite may be (probabilistically) produced. In contrast, in our model, community assembly occurs with the constraint that the initially supplied substrate can only be broken down into metabolites one or two steps below it in a free-energy hierarchy. This, combined with the embedded cellular proteome dynamics and thermodynamic constraints on free-energy availability on cellular growth rate, enforces a sequential and delayed differentiation of substrates during assembly time. This effectively introduces time lags between new substrates becoming available (see Fig 2C in the manuscript), leading to substantially longer transients during which rare species can establish on newly-generated resources (compare Fig 2A main text with Fig 1E in Marsland *et al.*). From this we find that the relationship between (free-) energy availability and steady state diversity is intrinsically linked to the process of ecosystem assembly. Low free-energy availability leads to a slower growth rate and thus a slower rate of substrate diversification (see Fig 4 main text). As we considered a top-down assembly procedure (one without additional immigration events), the number of niches (substrates) that are generated before species from the initially inoculated community reach extinction significantly determines the final diversity.

As previously mentioned, these key insights could only be gained due to the inclusion of both proteomic and thermodynamic constraints in our model. Firstly, the introduction of a (thermodynamically motivated) structure to the network of possible reactions increases the number of steps required for the system to fully develop. This means that the rate of ecosystem development becomes key. Secondly, our use of reversible enzyme kinetics provides a physically sound basis to determine the rate at which free-energy can be extracted from the (metabolite) environment. Thirdly, introducing proteomic constraints allows growth rates, for a particular rate of free-energy extraction, to be determined in a empirically supported manner. Taken together this allows our model to capture qualitatively how free-energy availability impacts cellular growth rate.

### S7 Interactions between functional groups

In the main text (see Fig 3), we assign species to functional groups based on the substrate of their most expressed reaction. It logically follows that competition interactions will mostly occur between members of the same functional group. In the case of facilitation a species breaks down a metabolite into a different metabolite which a second species breaks down. From this, we would expect facilitation interactions to (generally) occur between rather than within functional groups. By a similar logic, syntrophy interactions would also be expected to predominately occur within functional groups. Additionally, we would expect pollution interactions to frequently occur within functional groups as reactions in our model that share a substrate often share a waste-product. However, as waste-products can vary we would not expect this to be as exclusively the case as for competition interactions.

To test this hypothesis we produced heatmaps of both the total strength (Fig ST3) and frequency (Fig ST4) of the different interaction types across our simulations. Our general hypothesis holds with competition and pollution occurring generally within functional groups, and facilitation and syntrophy generally occurring between functional groups. This is observed regardless of whether the total strength or frequency of interactions is considered, though the pattern is considerably stronger for interaction strength. For the low-free energy condition we consider, comparatively few surviving species or functional groups are observed (see Figs 3 & 4 main text). The functional groups that survive are also likely to be those using substrates higher in the metabolite hierarchy, i.e. those that become available earlier in the process of sequential substrate diversification. For this reason, we only show the first 12 functional groups in our plots as the other 12 occur only extremely rarely. The same effect also leads to functional group 1 being by far the most prevalent functional group. This means that most facilitation and syntrophy interactions involve functional group 1 (see Figs ST3B-C and Figs ST4B-C). When considering the frequency of interactions this means that interactions within the first functional group are almost always the most common, regardless of the broader pattern (see Fig ST4). Finally, it is worth noting the mirroring that emerges between the facilitation and syntrophy. Regardless of whether total interaction strength (Figs ST3B-C) or frequency (Figs ST4B-C) is considered, facilitation is more prominent below the diagonal and syntrophy is more prominent above the diagonal, and there is a clear mirroring between the two interactions. Though the interaction strengths differ, each syntrophy interaction is clearly associated with a facilitation interaction.

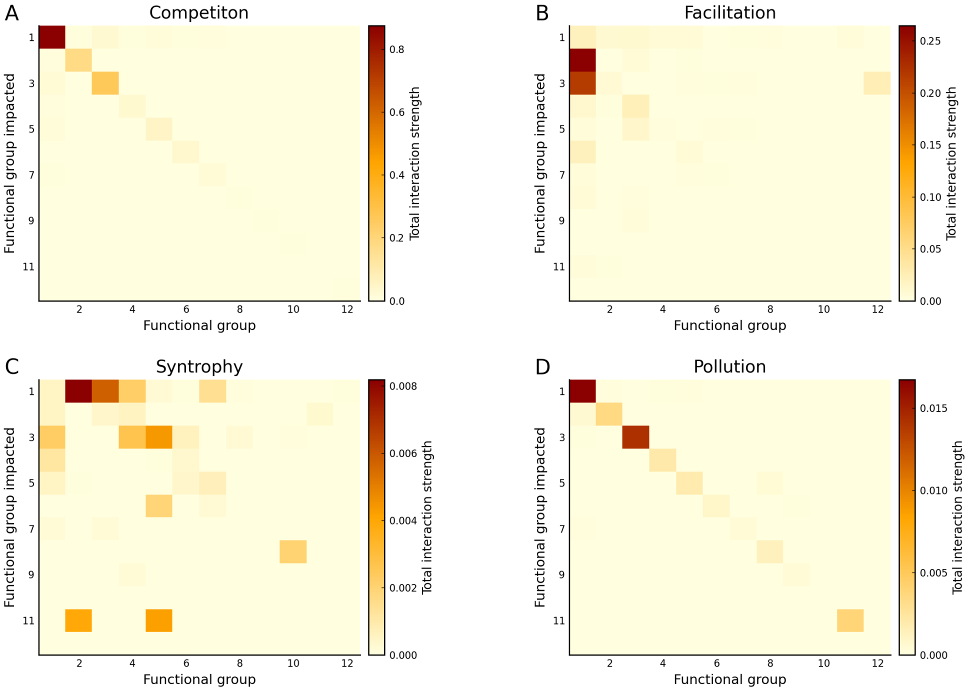

Figure ST3: **Heat maps of total interaction strength within and between functional groups.** The data were generated from our simulations for low-free energy (recalcitrant) substrates ($1.5 \times 10^6$ J mol$^{-1}$) with fully reversible kinetics. **A**: The total competition interaction strength is highest within functional groups, with the total strengths associated with interactions between different functional groups being negligible in almost every case. **B**: Total facilitation interaction strengths are relatively higher between functional groups compared to within functional groups. **C**: The relatively highest total interactions strengths are also between functional groups in the syntrophy case. **D**: In the case of pollution total interaction strengths are relatively highest within functional groups, with almost negligible total strengths between functional groups.

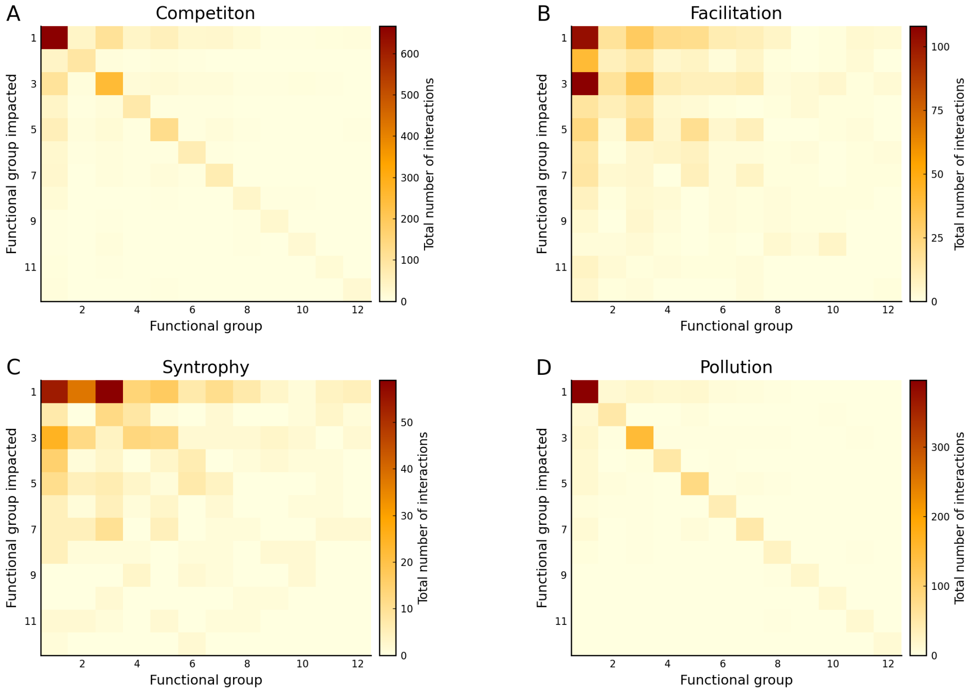

Figure ST4: **Heat maps of interaction frequency within and between functional groups.** The data were generated from our simulations for low-free energy (recalcitrant) substrates ($1.5\times10^6$ J mol$^{-1}$) with fully reversible kinetics. **A**: Almost all competition interactions are within the same functional groups. **B**: Though there are significant facilitation interactions within functional group 1, in general there are more interactions between functional groups. **C**: A similar pattern is observed for syntrophy with the majority of interactions being between functional groups. **D**: In the case of pollution interactions occur almost solely within functional groups.

#### S8 Model parameters

| | Symbol | Value | Units | Source |
|---|---|---|---|---|
| ATP free energy | $\Delta G_{\text{ATP}}$ | 75.0 | kJ mol$^{-1}$ | [6] |
| Temperature | $T$ | 293.15 | K | * |
| Cell mass | $m$ | $1 \times 10^8$ | # amino acids | [10] |
| Maximum elongation rate | $\gamma_m$ | 1260.0 | # steps min$^{-1}$ | [10] |
| Average ribosome mass | $n_R$ | 7459 | # amino acids | [7] |
| Average metabolic protein mass | $n_P$ | 300 | # amino acids | [8] |
| Rate of biomass loss | $d$ | $6 \times 10^{-5}$ | s$^{-1}$ | * |
| Elongation step cost | $\chi$ | 29 | # ATP | [13] |
| Elongation half saturation constant | $\gamma_{\frac{1}{2}}$ | $5 \times 10^8$ | # ATP cell$^{-1}$ | * |
| Average fraction of ribosomes bound | $f_b$ | 0.7 | N/A | [11] |
| Housekeeping proteome fraction | $\phi_Q$ | 0.45 | N/A | [12] |
| Ribosome fraction half saturation constant | $\Omega_{\frac{1}{2}}$ | $1 \times 10^9$ | # ATP cell$^{-1}$ | * |
| Metabolite dilution rate | $\rho$ | $6 \times 10^{-5}$ | s$^{-1}$ | [24] |
| Metabolite supply rate | $\kappa$ | $3.3 \times 10^{-7}$ | M s$^{-1}$ | * |
| Initial population | $N_0$ | 1.0 | cell | * |
| Initial metabolite concentration | $C_0$ | 0.0 | M | * |
| Initial intracellular energy concentration | $a_0$ | $1 \times 10^5$ | # ATP cell$^{-1}$ | * |
| Initial ribosome fraction | $\phi_{R,0}$ | 12.8 | % | * |

Table ST1: Table of fixed parameters in our model, kept constant across all simulated conditions. Parameters not taken from the literature are marked with an asterisk.

| | Symbol | Distribution | $\mu$ | $\sigma$ | Magnitude | Units |
|---|---|---|---|---|---|---|
| Maximum forward reaction rate | $k$ | Log-normal | 0 | $1/\ln 2$ | 10.0 | s$^{-1}$ |
| Substrate half saturation constant | $K_S$ | Log-normal | 0 | $1/\ln 2$ | $1.375 \times 10^{-3}$ | M |
| Reversibility factor | $r$ | Log-normal | 0 | $1/\ln 2$ | 10.0 | N/A |

Table ST2: Table of reaction kinetic parameters. These are drawn randomly for each reaction for each species, so the parameters of the random distributions used are given. The random numbers generated from these distributions are then multiplied by their respective magnitude in order to obtain final parameter values.

#### S9 Analysis of assumptions and parameterisations underlying our model

This section summarises all the key assumptions, implicit or explicit, underlying our model. We also address the choice of several key parameter values. Both of these things, along with our justifications for them, are contained in Tab ST3. In a few cases the justification for our assumptions goes beyond what can be contained in a table as indicated. These are our choice of values for the two saturation constants $\gamma_{\frac{1}{2}}$ and $\Omega_{\frac{1}{2}}$, the robustness of our results to changes in the parameterisation of the proteome allocation model, and the use of single-reactant single-product reactions. All of these are discussed in further detail in separate subsections.

| Assumption | Justification |
|---|---|
| Proteome can be partitioned into three compartments: ribosomes, metabolic proteins, and a fixed housekeeping fraction. | Assumption has been successfully used to model bacterial growth trade-offs, resulting in novel growth laws [1]. |
| Only catabolic reactions are directly included in our model. | Model aims to investigate the effect of free-energy supply on ecosystem dynamics, catabolism is more impacted by this than anabolism. |
| All metabolic protein can be treated as contributing to catabolism. | A portion of the metabolic protein fraction could be assigned to anabolism reducing the energy acquisition rate. However, this change could instead be absorbed into a reduction of the maximum enzyme rate ($k$). |
| Metabolic protein required to sustain catabolism does not vary between species. | In the model biomass synthesis pathways do not vary between species. This means a species with a catabolism requiring greater protein investment (e.g. one with more thermodynamic bottlenecks), would receive no benefit for the additional cost. |
| Ideal ribosome fraction (Eq 7 main text) is of saturating functional form. | Required in order to recover the empirically observed growth laws (see Fig ST1). |
| Effective elongation rate (Eq 6 main text) is of saturating functional form. | Maximum (per ribosome) translation rate has to be bounded. |
| Value of housekeeping proteome fraction ($\phi_Q$) is fixed across species. | As no compensating mechanism exists (e.g. chemotaxis or motility [28]), species with a higher housekeeping fraction would only ever be disadvantaged. |
| Fixed values are used for the two (energy) saturation constants $\gamma_{\frac{1}{2}}$ and $\Omega_{\frac{1}{2}}$. | More detail provided later in this section |

| | |
|---|---|
| Cells use free-energy solely for protein translation. | Translation is one of the most most energy consuming cellular processes [13]. Many other growth associated costs can reasonably be expected to scale with protein translation. Other uses of energy (e.g. motility) are beyond scope of the model. |
| The ATP cost per translation step $\chi$ does not vary between species or as cellular conditions change. | Simplifying assumption. As no benefit is attached to increased translation cost, allowing it to vary would simply disfavour strains with high $\chi$. |
| Model considers only single step enzyme reactions. | Simplifying assumption. However, more complex reactions can be captured by species possessing multiple reactions in series. |
| Reactions can only descend one or two steps downwards in the metabolite hierarchy. | Necessary as the model is intended to capture simple one-step enzyme reactions. |
| Fraction of ribosomes bound and translating ($f_b$) does not vary with changing cell conditions. | The value used is taken from the literature [11]. Increasing $f_b$ would lead to an increase in the growth rate, but would not change the overall results as this increased growth rate has to be met by an increased rate of free-energy acquisition. |
| Only considered single-reactant single-product reactions. | More detail provided later in this section. |
| Only external metabolites are modelled, e.g. no internal metabolite concentrations. | Reasonable first approximation. Thermodynamic limits will still apply as discussed in the Discussion section of the main text. |
| Constant temperature across all simulations. | The free-energy change caused by plausible temperature changes are very small compared to the variations in substrate free-energy we consider. |
| The metabolic protein mass ($n_p$) is the same for all reactions. | Simplifying assumption. Allowing this to vary would make some reactions less beneficial, but beyond reasonable scope of the paper to test this. |

| | |
|---|---|
| High free-energy change processes (e.g. protein translation) treated as irreversible. | Reversible and irreversible Michaelis–Menten kinetics are identical in the high dissipation (i.e. high free-energy change) limit. |
| Relative reaction expressions (see Eqs S15 & S16) vary between species but don't change over time. | Computationally intensive to implement, reaction expression for the whole community can still shift through species sorting. |
| The rate at which the proteome fraction can shift (see Eq S11) is set by a characteristic growth time scale $\tau_g$ (Eq S12). | Captures the fact that synthesised protein is retained for a significant period. $\tau_g$ is small enough that steady-state proteome fractions are reached. |
| Initial conditions ($N_0$, $C_0$, $a_0$, $\phi_{R,0}$) are the same across all simulations. | For all but extreme choices (which lead to all species going extinct) the choice of initial conditions does not effect the final steady states |
| Kinetic parameters ($k$, $K_S$, $r$) vary by approximately an order of magnitude around a fixed value. | Important to include kinetic variation. However, variation is kept relatively small to avoid kinetic variation dominating competition between strains. |
| For all simulations the same biomass loss rate ($d$) is used. | Changing this parameter alters the steady-state populations and the speed of ecosystem assembly. However, the increased population growth would be equal regardless of substrate free-energy and thus our results would still hold. |
| For all simulations the supply rate of the initial substrate $\kappa$ is the same. | This is an alternative way of changing the rate of free-energy supply into the system. We kept this fixed in order to investigate the impact of substrate free-energy. |

Table ST3: Table summarising the assumptions made in our model, with justifications given in each case. Cases where more justification is required are marked as "More detail provided later in this section".

#### S9.1 Choosing energy saturation constants

Our model contains two saturation constants that govern how cellular processes change as the internal energy concentration changes. They are the half-maximum constant for translational elongation rate $\gamma_{\frac{1}{2}}$ (see main text Eq 6), and the half-maximum constant for 'ideal' ribosome fraction $\Omega_{\frac{1}{2}}$ (see main text Eq 7). Both of these parameters are highly phenomenological; thus we cannot obtain values for them from the existing literature. We therefore had to pick reasonable values, chosen under the following restrictions: The first restriction arises from the fact that the two saturation constants must be of a similar order of magnitude for the proteome model to function. If $\gamma_{\frac{1}{2}}$ is substantially larger, then cells will end up with a large ribosome fraction that only operates at a small fraction of the maximum rate. Whereas if $\Omega_{\frac{1}{2}}$ is substantially larger, ribosomes will always operate at maximum speed but only a vanishingly small fraction of the proteome will be dedicated to them. Neither scenario is consistent with the empirically observed growth laws. The second restriction is that the magnitude of the two saturation constants cannot be too high or too low. In the high value limit, cell populations would never increase due to being unable to achieve sufficient translation rates or ribosome fraction to match the biomass loss rate ($d$). In the low value limit, cells would rapidly reach maximum translation rate and ribosome fraction, they would show very fast growth but would not be able to sustain the very high energy demand associated with this growth rate, leading to them exhausting their internal energy and dying out. This would of course be biologically unreasonable.

Based on these restrictions the saturation constants must be of intermediate value, and of similar order of magnitude to each other. We specifically chose $\Omega_{\frac{1}{2}}$ to be slightly larger than $\gamma_{\frac{1}{2}}$ to ensure that translation saturates quicker than the ribosome fraction shifts, in order to avoid the situation where species frequently fail to grow due to possessing a large number of inefficient ribosomes. Beyond this the specific values chosen are fairly arbitrary. However, we do not believe that changes in the values of these constants would effect the broad results of the paper, provided pathological cases are avoided by keeping to the restrictions set out in the prior paragraph. The two fundamental constraints on growth in our model are the availability of energy and the number of ribosomes available for protein translation. Increasing energy availability will thus always benefit the cells regardless of the choice of saturation constant values, provided a non-pathological choice is made. Thus, our broad results on the effect of (free) energy availability on the assembly of ecosystems would hold provided all strains are given the same values for the saturation constants.

The robustness of our results to changes in the values of the saturation constants is demonstrated in Fig ST5, where the original results for final number of surviving species is compared with those for three alternative parameterisations. These new parameterisations involved increasing the relative size of the translation rate half-maximum constant ($\gamma_{\frac{1}{2}}$) compared to the ribosome half-maximum constant ($\Omega_{\frac{1}{2}}$), increasing the value of both saturation constants, and decreasing the value of both saturation constants, respectively. In each case, our broad result that free-energy supply has the biggest impact on final diversity is recovered. Significant differences between enzyme schemes for the low free-energy supply case are not always recovered. However, this likely arises from the fact that less simulations were run for the robustness cases (50 simulations instead of 250), which lead to larger confidence intervals. Increasing both saturation constants resulted in significantly higher final diversities, and reducing both saturation constants parameterisation resulted in significantly lower final diversities. When species in our model possess higher saturation constants they grow more slowly, with more of the free-energy obtained being retained as ATP rather than used for growth. This slower growth and greater storage of ATP allows sub-optimal competitors for early substrates to survive in the system longer, which in turn allows some of them to take advantage of substrates that become available later, leading to increased final diversity.

##### S9.2 Robustness of our results to changes in proteome parameterisation

We now consider the robustness of our results to changes in the parametrisation of the proteome allocation model. To do this we consider three parameters. Firstly, we consider the average metabolic protein mass $n_P$. Though value of this parameter was estimated from the literature [8], increasing

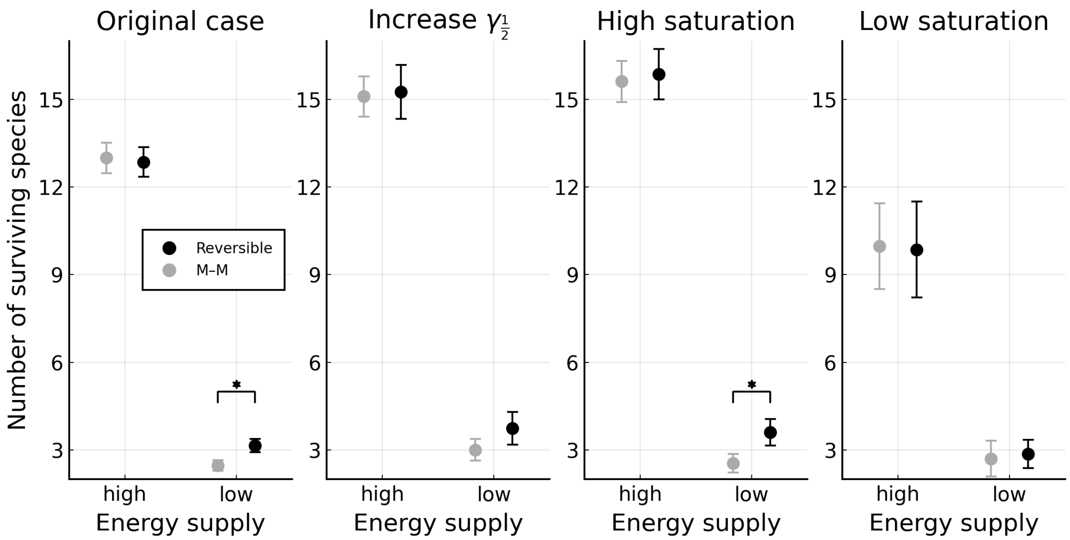

Figure ST5: **Effect of changing saturation constants.** Each panel shows a comparison of the final number of surviving strains between regimes (analogous to main text Fig 3B). For each regime, the average and 99% confidence intervals are plotted. For both energy regimes, we consider both reversible (black symbols) and Michaelis–Menten (M–M grey symbols) kinetics. Cases with a significant difference between these pairs are marked with a star ($P < 0.01$). The first panel shows the original result (averaged across 250 simulations, see Fig 3B), and the subsequent panels show the result for three alternative parameterisations (averaged across 50 simulations each). These alternative parameterisations are as follows: increased relative size for the translation rate half-maximum constant ($\gamma_{\frac{1}{2}} = 1 \times 10^9$ & $\Omega_{\frac{1}{2}} = 5 \times 10^8$), both saturation constants increased by a factor of 10 ($\gamma_{\frac{1}{2}} = 5 \times 10^9$ & $\Omega_{\frac{1}{2}} = 1 \times 10^{10}$), and both saturation constants decreased by a factor of 10 ($\gamma_{\frac{1}{2}} = 5 \times 10^7$ & $\Omega_{\frac{1}{2}} = 1 \times 10^8$).

its value offers a straightforward means of capturing the proteomic cost of anabolism, which is not explicitly included in our model. Secondly, we consider the average fraction of ribosomes bound $f_b$. This parameter was estimated from the literature [11], but would be expected to vary significantly between cells and across conditions. Finally, we consider the housing keeping proteome fraction $\phi_Q$. Again this fraction was estimated from the literature [12], but it would be expected to vary between species, particularly as it can include protein allocated to processes such as sensing and signalling [28]. The robustness of our results to changes in these three parameters is demonstrated in Fig ST6, where consistent with the previous robustness plot free-energy remains the largest driver of final species diversity, regardless of parameterisation. Final diversities decrease when the average metabolic protein mass increases, as the same investment in metabolic protein now results in less enzyme (see Eq 9 main text), leading to lower reaction rates. When the average fraction of ribosomes bound ($f_b$) is increased, the final diversities do not significantly change. Increasing the housekeeping fraction ($\phi_Q$), leads to a smaller proteome fraction to be split between ribosomes and metabolic protein thus decreasing the final diversities.

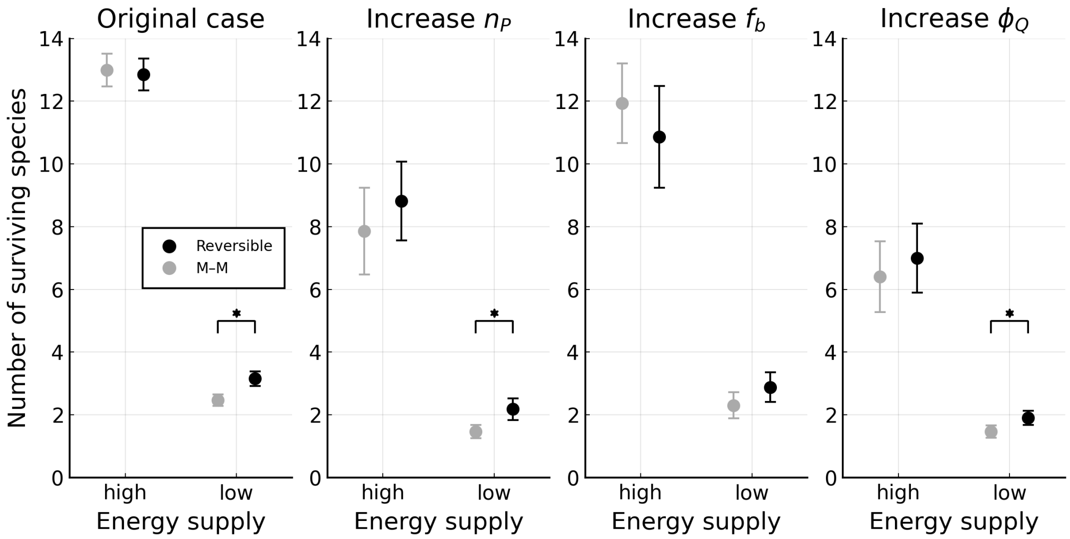

Figure ST6: **Effect of changing proteome parameters.** Each panel shows a comparison of the final number of surviving strains between regimes (analogous to main text Fig 3B). For each regime, the average and 99% confidence intervals are plotted. For both energy regimes, we consider both reversible (black symbols) and Michaelis–Menten (M–M grey symbols) kinetics. Cases with a significant difference between these pairs are marked with a star ($P < 0.01$). The first panel shows the original result (averaged across 250 simulations, see Fig 3B), and the subsequent panels show the result for three alternative parameterisations (averaged across 50 simulations each). These alternative parameterisations are as follows: increased average metabolic protein mass ($n_P = 600$), increased fraction of ribosomes bound ($f_b = 0.9$), and increased housekeeping proteome fraction ($\phi_Q = 0.7$).

##### S9.3 Assumption of single-reactant single-product reactions

In our model microbes can only make use of reactions that use a single reactant to produce a single product (with ATP production assumed to be coupled to this reaction). This represents a significant reduction in complexity from the realistic scenario of microbes generally possessing reactions involving many reactants resulting in many products. We made this simplification in order to reduce the analytic and computational complexity of the model, which made our model both more interpretable and easier to simulate. Additionally, we do not believe that including reactions involving multiple reactants or products would weaken the thermodynamic results that we obtained. In fact we expect that thermodynamic effects would in general be more significant when reactions using multiple reactants or products are included.

To see the thermodynamic effects we should note that in the multiple-reactant multiple-product case the reaction quotient (for a particular reaction $\alpha$) takes the form

$$Q(\boldsymbol{S}_\alpha, \boldsymbol{W}_\alpha) = \frac{\prod_{j=1}^{G_\alpha} (W_{j,\alpha})^{w_{j,\alpha}}}{\prod_{i=1}^{R_\alpha} (S_{i,\alpha})^{s_{i,\alpha}}}, \tag{S25}$$

where $W_{j,\alpha}$ is the concentration of the $j^{\text{th}}$ product, $S_{i,\alpha}$ is is the concentration of the $i^{\text{th}}$ reactant, $w_{j,\alpha}$ is the product coefficient of product $j$, $s_{i,\alpha}$ is the reactant coefficient of reactant $i$, $R_\alpha$ is the number of distinct reactants involved in the reaction and $G_\alpha$ is the number of distinct products involved in the reaction. As mentioned in the main text, in the single-reactant single product case this reduces to $Q(S_\alpha, W_\alpha) = W_\alpha / S_\alpha$. The value of the reaction quotient impacts the thermodynamic factor $\theta$ (Eq 11 main text), with higher values increasing the thermodynamic factor and lower values decreasing it. How close a reaction is to thermodynamic equilibrium is set by this thermodynamic factor, and therefore this is also (partly) determined by the value of the reaction quotient. When more metabolites are involved in the reaction the multiplicative form of the reaction quotient (see Eq S25) means that it varies more quickly with changing metabolite concentrations. So, while thermodynamic effects aren't guaranteed to be more important when considering reactions with multiple reactants and/or products, the variation in metabolite concentrations required to make these thermodynamic effects significant is smaller.

### References

1. Scott, M., Klumpp, S., Mateescu, E. M. & Hwa, T. Emergence of robust growth laws from optimal regulation of ribosome synthesis. *Molecular Systems Biology* **10,** 747–747. ISSN: 1744-4292 (Aug. 2014).
2. Bock, R. M. & Alberty, R. A. Studies of the Enzyme Fumarase. I. Kinetics and Equilibrium. *Journal of the American Chemical Society* **75,** 1921–1925. ISSN: 0002-7863, 1520-5126 (Apr. 1953).
3. Hoh, C. Y. & Cord-Ruwisch, R. A practical kinetic model that considers endproduct inhibition in anaerobic digestion processes by including the equilibrium constant. *Biotechnology and Bioengineering* **51,** 597–604. ISSN: 0006-3592 (Sept. 1996).
4. Hill, T. L. *Free energy transduction and biochemical cycle kinetics* ISBN: 978-1-4612-3558-3 (Springer-Verlag, New York, 1989).
5. Lynch, T., Wang, Y., van Brunt, B., Pacheco, D. & Janssen, P. Modelling thermodynamic feedback on the metabolism of hydrogenotrophic methanogens. *Journal of Theoretical Biology,* S0022519319302164. ISSN: 00225193 (May 2019).
6. Buckel, W. & Thauer, R. K. Energy conservation via electron bifurcating ferredoxin reduction and proton/Na+ translocating ferredoxin oxidation. *Biochimica et Biophysica Acta (BBA) - Bioenergetics* **1827,** 94–113. ISSN: 00052728 (Feb. 2013).
7. Keseler, I. M. *et al.* EcoCyc: a comprehensive database of Escherichia coli biology. *Nucleic Acids Research* **39,** D583–D590. ISSN: 0305-1048, 1362-4962 (Jan. 2011).
8. Brandt, F. *et al.* The Native 3D Organization of Bacterial Polysomes. *Cell* **136,** 261–271. ISSN: 00928674 (Jan. 2009).

9. Weiße, A. Y., Oyarzún, D. A., Danos, V. & Swain, P. S. Mechanistic links between cellular trade-offs, gene expression, and growth. *Proceedings of the National Academy of Sciences* **112,** E1038–E1047. ISSN: 0027-8424, 1091-6490 (Mar. 2015).

10. Bremer, H. & Dennis, P. in *Escherichia coli and Salmonella* (ed Neidhardt, C.) 1553–1569 (ASM Press, Washington, DC, 1996).

11. Underwood, K. A., Swartz, J. R. & Puglisi, J. D. Quantitative polysome analysis identifies limitations in bacterial cell-free protein synthesis. *Biotechnology and Bioengineering* **91,** 425–435. ISSN: 0006-3592, 1097-0290 (Aug. 2005).

12. Scott, M., Gunderson, C. W., Mateescu, E. M., Zhang, Z. & Hwa, T. Interdependence of Cell Growth and Gene Expression: Origins and Consequences. *Science* **330,** 1099–1102. ISSN: 0036-8075, 1095-9203 (Nov. 2010).

13. Lynch, M. & Marinov, G. K. The bioenergetic costs of a gene. *Proceedings of the National Academy of Sciences* **112,** 15690–15695. ISSN: 0027-8424, 1091-6490 (Dec. 2015).

14. Advani, M., Bunin, G. & Mehta, P. Statistical physics of community ecology: a cavity solution to MacArthur's consumer resource model. *Journal of Statistical Mechanics: Theory and Experiment* **2018,** 033406. ISSN: 1742-5468 (2018).

15. Cui, W., Marsland, R. & Mehta, P. Effect of Resource Dynamics on Species Packing in Diverse Ecosystems. *Physical Review Letters* **125,** 048101. ISSN: 0031-9007, 1079-7114 (July 2020).

16. Levin, S. A. Community Equilibria and Stability, and an Extension of the Competitive Exclusion Principle. *The American Naturalist* **104,** 413–423. ISSN: 0003-0147, 1537-5323 (Sept. 1970).

17. Hutchinson, G. E. The Paradox of the Plankton. *The American Naturalist* **95,** 137–145. ISSN: 0003-0147, 1537-5323 (May 1961).

18. Huisman, J. & Weissing, F. J. Biodiversity of plankton by species oscillations and chaos. *Nature* **402,** 407–410. ISSN: 0028-0836, 1476-4687 (Nov. 1999).

19. Descamps-Julien, B. & Gonzalez, A. Stable coexistence in a fluctuating environment: an experimental demonstration. *Ecology* **86,** 2815–2824. ISSN: 0012-9658 (Oct. 2005).

20. Huisman, J., van Oostveen, P. & Weissing, F. J. Species Dynamics in Phytoplankton Blooms: Incomplete Mixing and Competition for Light. *The American Naturalist* **154,** 46–68. ISSN: 0003-0147, 1537-5323 (July 1999).

21. Roughgarden, J. & Feldman, M. Species packing and predation pressure. *Ecology* **56,** 489–492. ISSN: 00129658 (Mar. 1975).

22. Yamamichi, M. & Letten, A. D. Rapid evolution promotes fluctuation-dependent species coexistence. *Ecology Letters* **24** (ed Adler, F.) 812–818. ISSN: 1461-023X, 1461-0248 (Apr. 2021).

23. Großkopf, T. & Soyer, O. S. Microbial diversity arising from thermodynamic constraints. *The ISME Journal* **10,** 2725. ISSN: 1751-7370 (Nov. 2016).

24. Posfai, A., Taillefumier, T. & Wingreen, N. S. Metabolic Trade-Offs Promote Diversity in a Model Ecosystem. *Physical Review Letters* **118,** 028103. ISSN: 1079-7114 (Jan. 2017).

25. Li, Z. *et al.* Modeling microbial metabolic trade-offs in a chemostat. *PLOS Computational Biology* **16** (ed Grilli, J.) e1008156. ISSN: 1553-7358 (Aug. 2020).

26. Pacciani-Mori, L., Suweis, S., Maritan, A. & Giometto, A. Constrained proteome allocation affects coexistence in models of competitive microbial communities. *The ISME Journal* **15,** 1458–1477. ISSN: 1751-7362, 1751-7370 (May 2021).

27. Marsland, R. *et al.* Available energy fluxes drive a transition in the diversity, stability, and functional structure of microbial communities. *PLOS Computational Biology* **15** (ed Morozov, A. V.) e1006793. ISSN: 1553-7358 (Feb. 2019).

28. Ni, B., Colin, R., Link, H., Endres, R. G. & Sourjik, V. Growth-rate dependent resource investment in bacterial motile behavior quantitatively follows potential benefit of chemotaxis. *Proceedings of the National Academy of Sciences* **117,** 595–601. ISSN: 0027-8424, 1091-6490 (Jan. 2020).
